# A Large Language Model–Powered Map of Metabolomics Research

**DOI:** 10.1101/2025.03.18.643696

**Authors:** Olatomiwa O. Bifarin, Varun S. Yelluru, Aditya Simhadri, Facundo M. Fernández

## Abstract

We present a comprehensive map of the metabolomics research landscape, synthesizing insights from over 80,000 publications. Using PubMedBERT, we transformed abstracts into 768-dimensional embeddings that capture the nuanced thematic structure of the field. Dimensionality reduction with t-SNE revealed distinct clusters corresponding to key domains such as analytical chemistry, plant biology, pharmacology, and clinical diagnostics. In addition, a neural topic modeling pipeline refined with GPT-4o mini reclassified the corpus into 20 distinct topics—ranging from “Plant Stress Response Mechanisms” and “NMR Spectroscopy Innovations” to “COVID-19 Metabolomic and Immune Responses.” Temporal analyses further highlight trends including the rise of deep learning methods post-2015 and a continued focus on biomarker discovery. Integration of metadata such as publication statistics and sample sizes provide additional context to these evolving research dynamics. An interactive web application (https://metascape.streamlit.app/) enables dynamic exploration of these insights. Overall, this study offers a robust framework for literature synthesis that empowers researchers, clinicians, and policymakers to identify emerging research trajectories and address critical challenges in metabolomics, while also sharing our perspectives on key trends shaping the field.

## Introduction

Metabolomics, the field that studies small molecules in biological systems has witnessed rapid growth over the past two decades since its inception in the late 1990s.^1^ This explosion is driven by advances in analytical instrumentation, big data analysis, and increasing recognition of the profound influence of metabolites on organismal biology.^2, 3^ Yet, as publication counts surge and research topics diversify—from plant metabolism^4^ to human disease biomarkers—^5^ it becomes increasingly difficult for researchers to track the broadening scope of work, identify emergent themes, and discover interdisciplinary connections. Traditional literature surveys and keyword-based database queries offer only partial insights, focusing on specific research questions or narrow subfields. While these targeted approaches are useful for in-depth analyses, they are less effective at painting a panoramic picture of the research landscapes. As a result, critical patterns—such as how computational methods are adopted over time, how studies cluster by organism or disease, and how novel techniques propagate across different disciplines—can remain hidden. Adapting the work of González-Márquez and colleagues on the landscape of biomedical research,^6^ in this work, we present a global framework for mapping the metabolomics literature in a two-dimensional “map”. Leveraging PubMedBERT,^7^ a domain-specific language model trained on PubMed data, we transform metabolomics-related publications into high-dimensional embeddings. We then apply dimensionality reduction methods, including t-Distributed Stochastic Neighbor Embedding (t-SNE) and Uniform Manifold Approximation and Projection (UMAP), to generate two-dimensional layouts that capture the global structure of the embeddings. To enhance the quality of embedding interpretation, in addition to inferring the category of paper abstracts from the titles of published journals, as previously demonstrated,^6^ we conducted topic modeling using a Large Language Model (LLM) – GPT-4o Mini from OpenAI – for improved distilled representation. This combination of natural language processing, LLM and manifold learning enables a comprehensive perspective on the field, highlighting how subdisciplines and research themes align or diverge. Our approach not only confirms the prominent role of analytical chemistry and clinical studies in metabolomics but also reveals the extent to which interdisciplinary journals and diverse applications shape the research corpus. To further enrich these observations, we incorporate metadata on study designs, sample sizes, and keyword queries, allowing for fine-grained exploration of publication characteristics. The resulting visualization provides an intuitive interface for identifying high-density research clusters—such as COVID-19–related metabolomics—and for tracking how emerging areas, like deep learning, are gaining traction over time. Additionally, to democratize access to our findings and empower broader discovery, we have developed an interactive web application (https://metascape.streamlit.app/). This platform allows users to filter and explore the map by keywords, authors, and research domains. By offering a global perspective on metabolomics, our work aims to support researchers, clinicians, and policymakers in navigating an ever-expanding literature, fostering deeper insights into our collective study of the biochemical underpinnings of life. Beyond metabolomics, this approach serves as a model for large-scale literature analysis in other fast-evolving disciplines.

## Mapping the Metabolomics Landscape: Empirical Insights and Trends

### Publication Analytics

The field of metabolomics has experienced exponential growth since its inception, with publication counts steadily increasing since 1998, to approximately 12,000 publications in 2023 in our dataset (**Figure 1a**). The lowest rate of growth occurred between 1998 and 1999, as expected during the early stages of the field’s emergence. In contrast, the highest rate of growth was observed between 2020 and 2021, likely influenced by the COVID-19 pandemic (**Table S1**). The mean annual growth rate across all year pairs was approximately 448 publications. Between 1998 and 2002, the field saw consistently low growth rates, reflecting its nascent stage, while the years 2011 to 2021 exhibited consistently higher growth, highlighting the maturation and broader adoption of metabolomics techniques (**Table S2**). An analysis of abstract lengths (**Figure 1b**) shows that most abstracts contain between 150 and 250 words, with a modal length of around 200 words, consistent with standard scientific publication practices and journal word limits. A long tail of shorter and longer abstracts likely reflects the varying requirements of different publication venues. Journal frequency analysis (**Figure 1c**) reveals that *Scientific Reports* and *Metabolites* are the leading journals, each publishing over 2,000 papers on metabolomics. Other notable journals include *PLOS ONE*, *Analytical Chemistry*, and the *International Journal of Molecular Sciences*. Additionally, a word cloud generated from the abstracts (**Figure 1d**) highlights key terms such as “mass spectrometry,” “identified,” “amino acid,” and “associated,” reflecting the field’s focus on metabolite identification, biochemical pathways, and quantitative techniques. The frequent use of terms such as “treatment” and “effect” underscores the clinical and applied dimensions of metabolomics, with many studies investigating the impacts of interventions or treatments on metabolic profiles. Less obvious keywords like “N” and “P” indicating number of samples and p value, respectively, are also observed in the word cloud.

**Figure 1.**
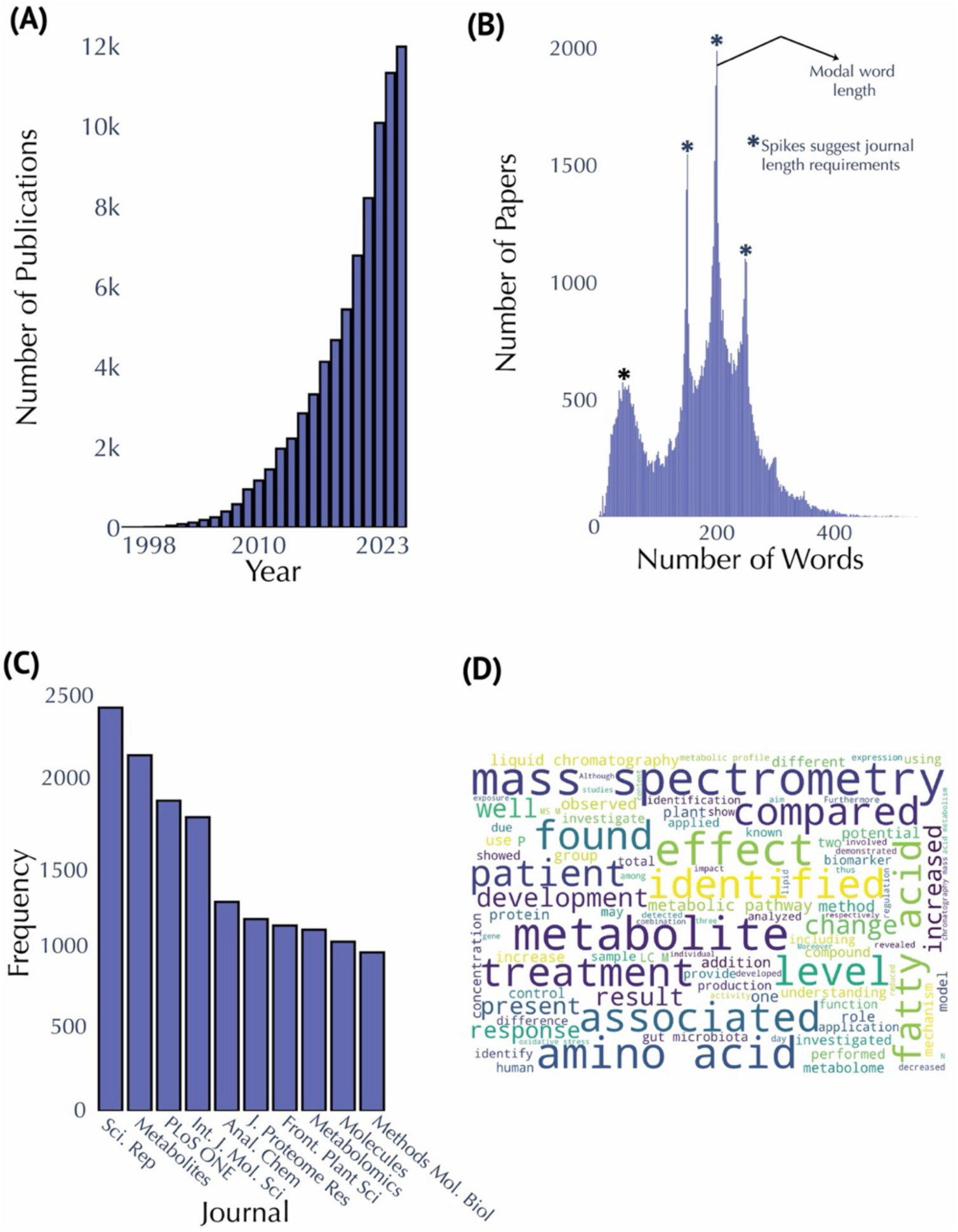
Publication and Abstract Analysis in Metabolomics Research. **(A)** Annual publication counts from 1998 to 2023. **(B)** Distribution of abstract lengths, highlighting a modal range of 150–250 words. Spikes suggest journal word limits. **(C)** Top ten publishing journals in metabolomics research, with *Scientific Reports* and *Metabolites* leading the field. **(D)** Word cloud generated from abstracts, larger sized text indicates increased frequency of occurrence.

### Global View of Metabolomics Corpus

We used a domain-specific language model, PubMedBERT^7^—trained on PubMed abstracts and full-text articles—to generate 768-dimensional embedding vectors for each publication abstract. To visualize these high-dimensional embeddings, we applied two dimensionality reduction techniques: Uniform Manifold Approximation and Projection (UMAP) and t-Distributed Stochastic Neighbor Embedding (t-SNE). Both methods reduced the 768-dimensional vectors to two dimensions, providing a global perspective on how publications cluster by research domain (**Figure 2a**, **Figure S1**). To evaluate each technique’s performance, we compared k-Nearest Neighbors (k-NN) accuracy and recall (**Table 1**). t-SNE outperformed UMAP on both metrics (k-NN accuracy: 0.56 vs. 0.54; k-NN recall: 0.33 vs. 0.14), indicating better preservation of the global embeddings in the local embedding structure. For additional context, we color-coded the reduced embeddings according to 18 predefined research categories inferred from the journal venues they are published (see **Methods** for details). Analytical chemistry emerged as the most represented field (8,981 publications), followed by plant biology (4,648) and pharmacology (4,229), while nephrology (277) and sports science & medicine (76) had fewer publications (**Table S3**). The t-SNE visualization (**Figure 2a**) reveals distinct clusters corresponding to major research areas, such as plant biology, analytical chemistry, environmental sciences, animal sciences, pharmacology, cancer biology, and immunology. Plant biology exhibits a compact cluster, suggesting strong thematic focus (**Figure 2b**), whereas the broader spread of analytical chemistry indicates wide applicability across metabolomics (**Figure 2c**). Overlapping clusters—like toxicology and environmental sciences (**Figure 2d-e**)—highlight interdisciplinary connections, which are similarly evident for other domains like cancer research and immunology, as well as animal science and food science. To broaden our “global view” of metabolomics publications beyond the two-dimensional t-SNE (**Figure 2a**) and UMAP (**Figure S1**) embeddings used for domain clustering, we examined temporal patterns across the corpus. We color-coded the t-SNE embeddings by publication year (**Figure S2**), revealing how early work (pre-2010) clustered primarily around analytical chemistry and plant biology. We then divided the dataset into eight discrete time periods (1998–2000 through 2022–early 2024) and generated separate t-SNE plots for each (**Figure S3**). Early clusters (1998–2009) are comparatively sparse and heavily oriented toward analytical methods, reflecting that analytical chemistry had the highest publication count before 2010, followed by plant biology (**Table S4**). In contrast, later periods exhibit denser, more diverse clusters, mirroring both the exponential growth of metabolomics and its expanding translational applications.

**Figure 2.**
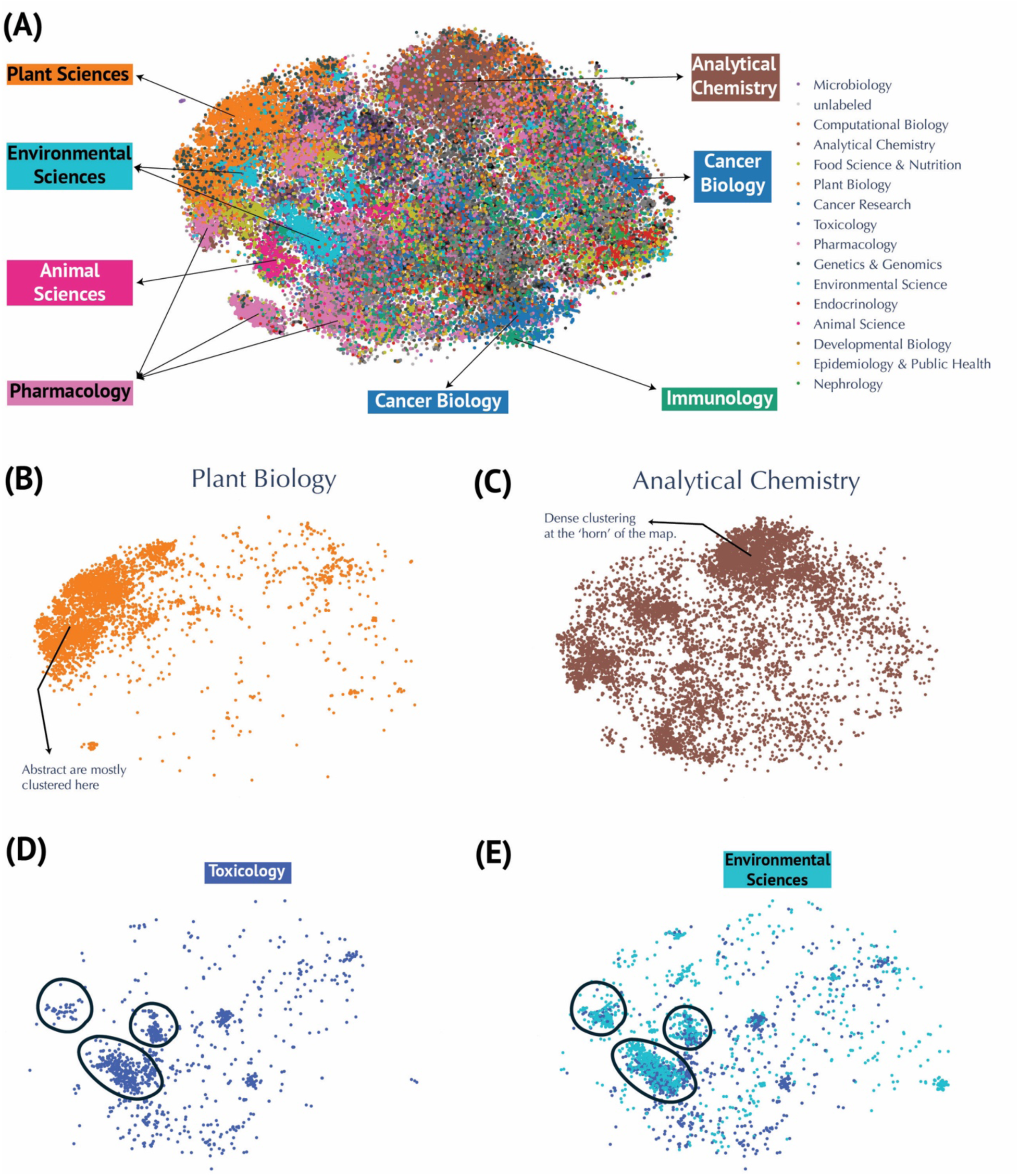
Visualization of Metabolomics Research Fields using t-SNE Embeddings. **(A)** Annotated t-SNE projection highlighting research domains. **(B)** Focused visualizations of clusters for Plant Biology and **(C)** Analytical Chemistry. Dense clustering is seen at the ‘horn’ of the map. (**D-E**) Examples of overlapping clusters, such as Environmental Sciences and Toxicology, illustrating connections on the map between related domains.

**Table 1.**
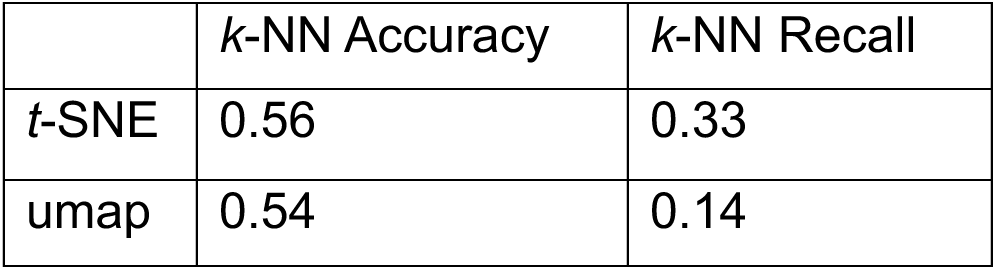
Performance evaluation of dimensionality reduction techniques. Comparison of t-SNE and UMAP on preserving the structure of 768-dimensional PubMedBERT embeddings in a two-dimensional space. Metrics include k-Nearest Neighbors (k-NN) accuracy and k-NN recall.

One limitation of inferring a publication’s research domain from the title of the journal is that many articles in multidisciplinary journals go unlabeled. Indeed, the “unlabeled” cluster comprises 41,721 publications (**Figure S5**; **Table S3**) and is broadly dispersed, reflecting the heterogeneity of papers that could not be assigned to predefined categories. This wide distribution highlights how multidisciplinary or non-specialized journals nevertheless contribute significantly to metabolomics research. To address this issue, we performed topic modeling using cTF–IDF for topic representation and GPT-4o mini for topic finetuning, thereby generating thematic labels that capture each cluster’s topical scope rather than merely its research field. For instance, in the cluster representing plant biology, the cTF–IDF computation returned keywords such as *plants*, *genes*, *compounds*, *stress*, *growth*, *species*, *leaves*, *biosynthesis*, and *production* (**Figure 3**). These words, together with representative abstracts, were provided to GPT-4o mini to produce more descriptive labels (see **Methods** for details). Overall, this approach revealed 20 distinct topics, including “Plant Stress Response Mechanisms,” “Metabolic Profiles and Dysregulation,” “Cancer Metabolism and Therapy Resistance,” “Metabolomics Data Analysis and Integration,” “Gut Microbiota and Metabolomic Interactions,” “Metabolomics in Neurodegenerative Disorders,” “Environmental Toxicology and Metabolism,” “Metabolomics in Animal Nutrition,” “Microbiota– Gut–Brain Axis Interactions,” “Kidney Disease Metabolomics and Biomarkers,” “Metabolomics in Rheumatoid Arthritis,” “Maternal Metabolomic Changes in Pregnancy,” “Lung Disease and Metabolic Dynamics,” “NMR Spectroscopy Innovations in Metabolomics,” “COVID-19 Metabolomic and Immune Responses,” “Lipid Profiling Techniques,” “Host–Parasite Metabolic Interactions,” “Male Fertility and Reproductive Metabolomics,” “Ocular Metabolomics and Disease Mechanisms,” and “Salivary Metabolomics in Oral Health” (**Figure 3**; **Figure 4a**). In contrast to the research field embeddings, which labeled ∼40,000 publications (**Figure S3**), the topic modeling assigned 65,676 publications to a defined topic (**Table S5**). Notably, the plant biology cluster (“Plant Stress Response Mechanisms”) remains compact (**Figure 4b**), consistent with its presentation in the research field embeddings (**Figure 2b**) and contains the most assigned publications at 19,258 (**Table S5**). The next largest topic, “Metabolic Profiles and Dysregulation,” encompasses various forms of altered metabolism in human diseases, reflected by discriminating keywords such as *liver*, *diabetes*, *insulin*, *muscle*, *mice*, *exercise*, *obesity*, *heart*, *diet*, and *hepatic*. (**Figure 3**) Meanwhile, “Cancer Metabolism and Therapy Resistance” appear with scattered clusters throughout the atlas, emphasizing its interdisciplinary nature (**Figure 4c**). The dense analytical chemistry cluster at the “horn” of the atlas (**Figure 2b**) is now partly decomposed into “Metabolomics Data Analysis and Integration” (**Figure 4d**) and “NMR Spectroscopy Innovations” (**Figure 4e**), most likely indicative of methods development.

**Figure 3.**
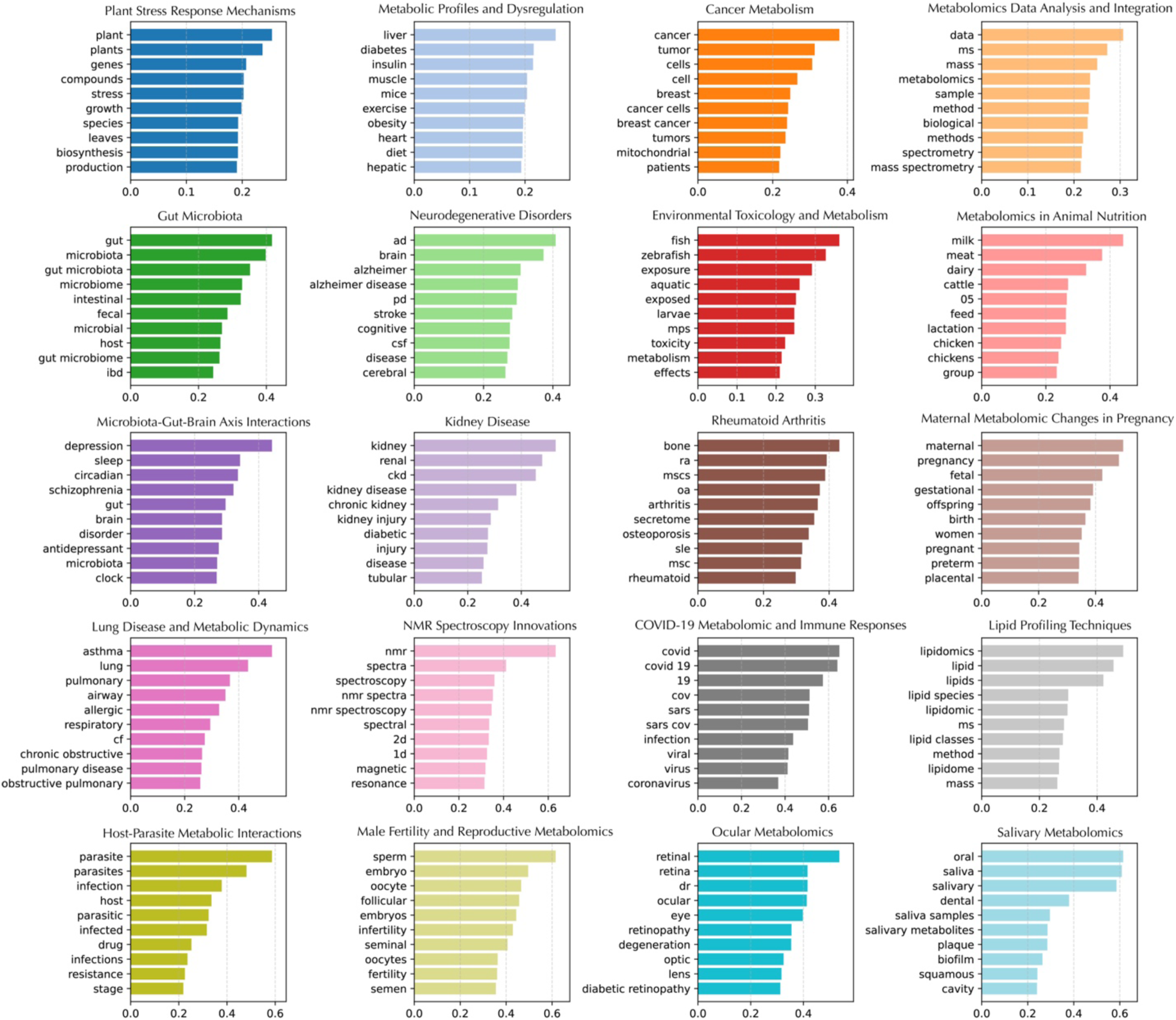
c-TF-IDF Topic Word Scores for Metabolomics Corpus. Each subplot represents a topic identified through BERTopic modeling of metabolomics literature. The x-axis shows the c-TF-IDF score, representing the importance of each word within the topic. The y-axis lists the top 10 most relevant words for each topic.

**Figure 4.**
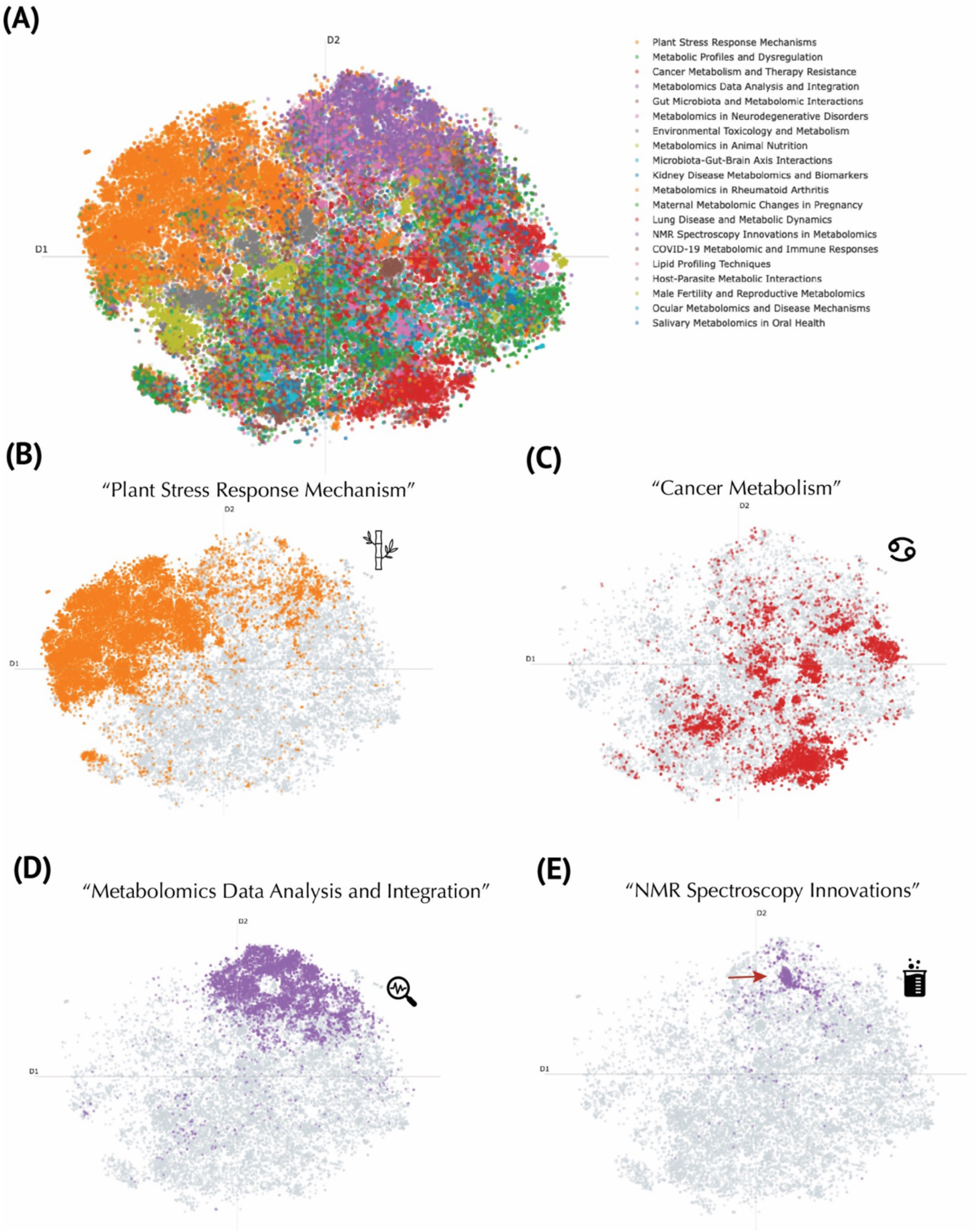
Topic Modeling of the Metabolomics Corpus Using a Large Language Model. **(A)** Global t-SNE projection of publications, colored by representative topics discovered through the BERTopic pipeline. The legend (top right) lists exemplar thematic labels generated by GPT4o mini. **(B)** Focus on publications labeled as “Plant Stress Response Mechanism” (orange). **(C)** Cluster of “Cancer Metabolism” publications (red), illustrating the concentration of work on metabolic pathways implicated in oncogenesis and therapy resistance. **(D)** “Metabolomics Data Analysis and Integration” topic (purple), marking computationally oriented publications. **(E)** “NMR Spectroscopy Innovations” topic (purple), distinct from the data-integration cluster, focused on methodological advancements in nuclear magnetic resonance techniques.

### Research Trend Discovery

We leveraged our global embedding space to pinpoint emergent topics and patterns in the metabolomics literature (**Figures 5 and 6**). Keyword-based queries proved particularly revealing. For example, studies whose abstracts include the phrase “for the first time” are dispersed throughout the map, signifying a diverse range of novel contributions (**Figure 5a**). By contrast, research on “COVID-19 or SARS-CoV-2” clusters densely in one region (**Figure 5b**), highlighting the concentrated surge in metabolomics investigations related to pandemic-driven questions, with publications from 2020 onward (**Figure S5**). Inspection of additional metadata features offered further insights into study designs and analytical practices. P-values are more prevalent in disease-focused or animal science contexts (**Figure 5c**), suggesting that hypothesis-driven experimental designs are particularly common in these domains. **Figure 5d** shows reported sample sizes concentrated in clinical and animal studies, with most samples under 50 (n = 1849; **Figure 5e**). Larger sample sizes—exceeding 1000— was only 51 (**Figure 5e**) and tends to appear in systematic meta-reviews. **Figures 6a–d** illustrate how searches for “biomarker discovery” and for “(pathway analysis OR metabolic pathway) AND mechanism” in the abstract occupy distinct but partially overlapping regions of the corpus. Biomarker-focused papers are widely distributed in the analytical chemistry and disease-related clusters, aligning with the mid-to-late 2000s rise in translational research, likely reflective of early publication efforts in biomarker discovery (**Figures 6c–d**). In contrast, mechanistic studies are particularly prominent in plant biology and pharmacology (**Figure 6a**) and skew slightly more recent (**Figure 6b**), suggesting ongoing exploration of metabolic pathways in these fields. We also contrasted classical multivariate approaches with newer machine-learning methods. While “chemometrics or multivariate analysis” is well-represented across multiple decades, abstracts mentioning “deep learning or neural network” show a comparatively late surge (**Figures 6e–f**). A more focused query on “deep learning” alone (**Figures 6g–h**) reveals that although still sparse, these publications cluster in data-science–oriented regions of the map and predominantly appear after 2015. Finally, we supplemented these findings by color-coding articles according to collaboration size, journal title length, and abstract length (**Figure S6**). Collaborative efforts (**Figure S6a**) range from small teams to large consortia, underscoring the varied scale of research. Journal title length (**Figure S6b**) does not appear to correlate with any particular domain, whereas the distinct pattern seen in the abstract length (**Figure S6c**) likely reflects specific journal guidelines and article types. Overall, this multifaceted embedding-based analysis offers a robust framework for charting how specific research foci and methodological approaches have emerged and evolved across the metabolomics literature.

**Figure 5.**
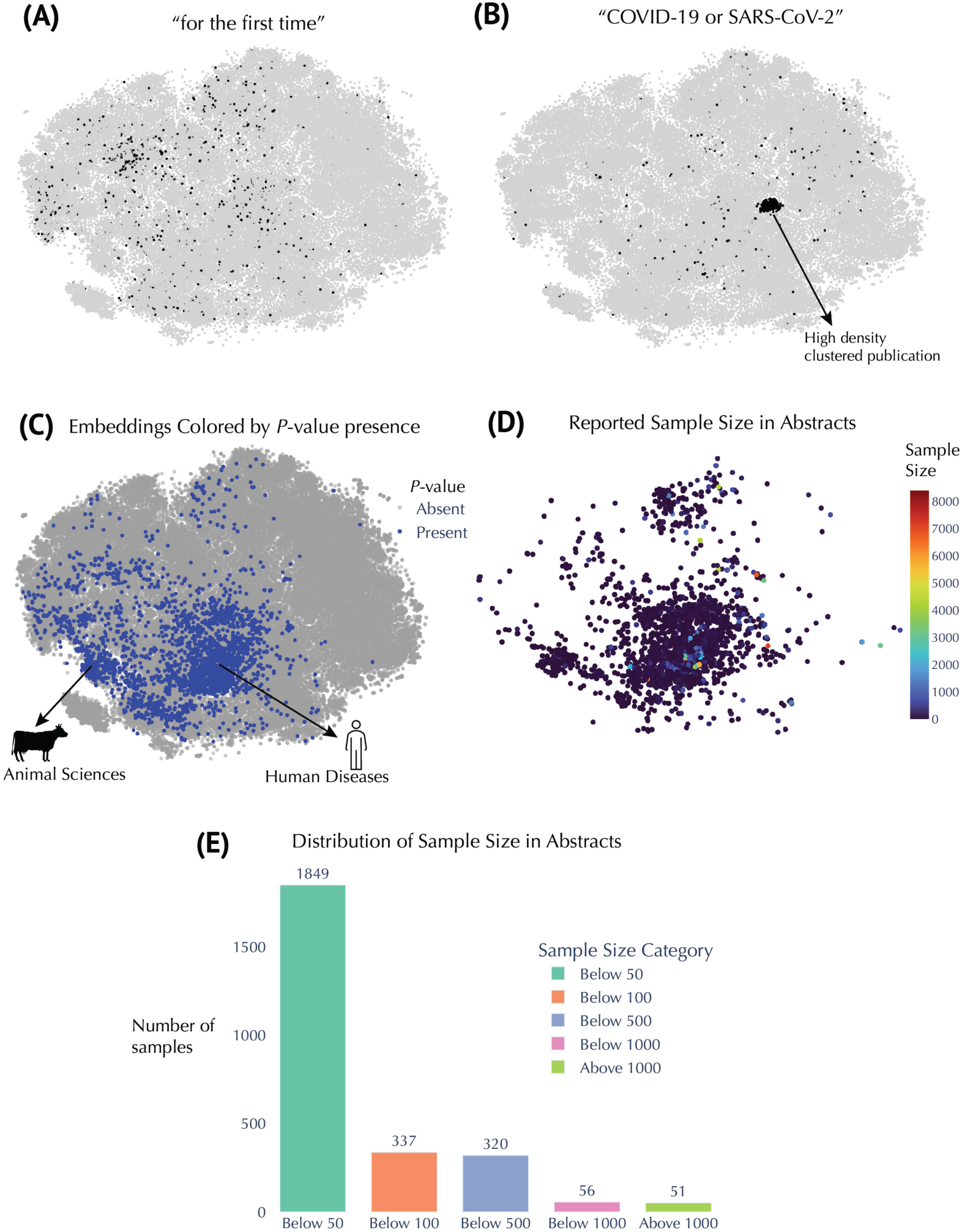
Insights from Embedding Metadata in Metabolomics Research. (**A**) t-SNE embeddings filtered to show abstracts containing the phrase *“for the first time.”* (**B**) t-SNE embeddings filtered for abstracts containing “COVID-19” or “SARS-CoV-2” (regex: COVID-19|SARS-CoV-2). (**C**) Embeddings colored by presence or absence of P-values, identified by the regex [pP]\s?[<=>]. For example, phrases such as “p<0.05,” “P=0.01,” or “p > 0.001” match this pattern. (**D**) Embeddings colored by reported sample size, extracted from expressions matching n\s?=\s?(\d+). For example, “n=30,” “n = 500,” or “n=1000” would be captured. (**E**) Distribution of these extracted sample sizes across the relevant abstracts.

**Figure 6.**
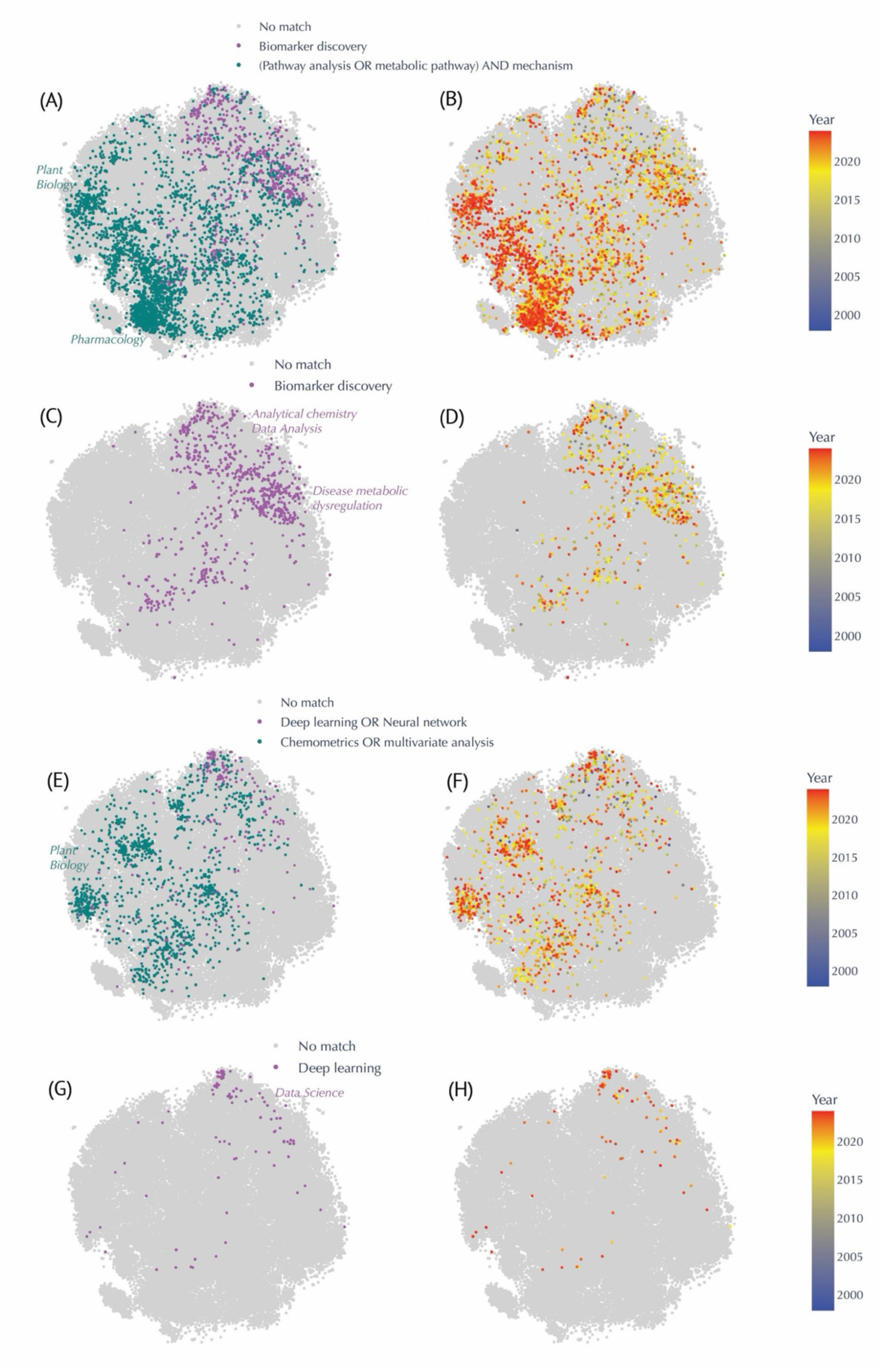
Evolution of Keyword-based Queries in the Metabolomics Corpus, Highlighting Shifts in Methodological Approaches and Conceptual Focuses Over Time. In each keyword panel, matched abstracts appear in color, while unmatched entries appear in light gray. The heatmap on the right panels transitions from **blue** (older publications) to **red** (more recent publications). **(A–B)** Comparison between “Biomarker discovery” (**teal**) and “(Pathway analysis OR metabolic pathway) AND mechanism” (**purple**). Papers mentioning “pathway mechanisms” cluster prominently in plant biology and pharmacology regions and are more recent (B), while “biomarker discovery” studies stretch across analytical chemistry and disease-related areas from the mid-2000s onward (A). **(C–D)** Focus on “Biomarker discovery” alone (purple) and its temporal distribution, revealing a surge in clinical and disease-oriented research in the late 2000s (D). **(E–F)** Contrast between “Deep learning OR neural network” (**purple**) and “Chemometrics OR multivariate analysis” (**teal**). Classical multivariate methods span earlier periods, whereas deep learning publications have intensified mainly post-2015 (F). **(G–H)** Spotlight on “Deep learning” (purple), showing relatively sparse coverage overall, yet relatively denser clustering in data science–focused areas in more recent years (H).

### Synthesis of Findings and Perspectives

Here, we present a comprehensive global map of metabolomics research by integrating advanced natural language processing with dimensionality reduction and topic modeling techniques. By leveraging PubMedBERT embeddings^7^, t-SNE, and BERTopic-based clustering,^8^ our methodology systematically delineates the thematic and temporal evolution of the metabolomics literature spanning 1998 to early 2024. This methodology not only enables the identification of key research domains but also captures the dynamic interplay of topics that underlie the progression of the field, thereby offering critical insights that extend beyond traditional literature reviews.

One notable finding is the sustained period of growth that began in 2011. Our temporal analysis reveals that the field experienced a marked acceleration in publication output starting that year. This acceleration may be attributed to factors such as heightened interest, increased funding, technological advancements, and the exploration of broader applications.^2^ Indeed, our map (**Figure S3**) demonstrates that metabolomics applications expanded significantly after 2010. Overall, this growth underscores both the maturation of metabolomics and its expanding influence across diverse domains, including analytical chemistry, clinical diagnostics,^9^ and plant biology.^10^

One unsurprising finding is the prominent role of analytical chemistry within metabolomics research. Our analyses reveal that analytical chemistry remains the dominant theme, as evidenced by its extensive representation in the corpus and the widespread distribution of its associated publications across the embedding space (**Figure 2c**). This reflects the foundational role that sophisticated analytical methods, such as mass spectrometry^11^ and NMR spectroscopy,^12^ play in the identification and quantitation of metabolites. In contrast, the plant biology cluster exhibits a more compact and tightly focused grouping, indicating a high degree of thematic cohesion in studies centered on plant stress responses and biosynthetic pathways^10^ (**Figure 2b**).

Our comparison of dimensionality reduction techniques also yielded important methodological insights, as also seen in González-Márquez et al.^6^ The performance evaluation, in which t-SNE demonstrated superior k-NN accuracy and recall compared to UMAP, highlights the utility of t-SNE in preserving local data structures when reducing high-dimensional embeddings to two dimensions (**Table 1**). This finding is critical because the fidelity of local neighborhood preservation directly impacts the interpretability of clusters and, by extension, the reliability of downstream topic modeling and metadata analyses.

A key strength of our approach is the integration of topic modeling with refined language generation. By applying a c-TF-IDF framework^8^ in conjunction with GPT-4o mini, a large language model from OpenAI, we were able to generate nuanced and informative labels for each cluster, thereby moving beyond the limitations of inferring research domains solely from journal titles^13^ (**Figure 2a & 4a**). This approach allowed for the reclassification of over 65,000 publications into 20 distinct topics, covering a wide range of research areas—from “Plant Stress Response Mechanisms” (**Figure 4b**) to “NMR Spectroscopy Innovations” (**Figure 4c**) and “COVID-19 Metabolomic and Immune Responses”. The detailed topic labeling not only facilitates a deeper understanding of thematic trends but also aids researchers in quickly identifying areas of novel contribution and emerging interest.

Furthermore, by leveraging keyword-based queries in abstracts, we were able to observe differences between two major thematic streams in metabolomics research: biomarker discovery and pathway mechanism elucidation. Biomarker-focused studies, predominantly found in analytical chemistry and clinical clusters, echo early translational efforts from the mid-2000s^14^ (**Figure 6c-d**). In contrast, mechanistic investigations, which are particularly prominent in plant biology and pharmacology, offer deeper insights into the biochemical pathways driving metabolic processes^15^ (**Figure 6a**). In addition, recent surge in studies employing deep learning for metabolomics data analysis were observed^16^ (**Figure 6g-h**).

Despite the impressive volume of research, a significant challenge in metabolomics is the low number of samples in many studies (**Figure 5d-e**), which carries important consequences. For instance, while numerous investigations report promising metabolic biomarkers, there remains a scarcity of clinically validated and widely adopted biomarkers.^17^ This gap is partly attributable to issues such as small sample sizes and insufficient validation across independent cohorts. For example, a meta-analysis on pancreatic cancer found that 87% of 655 potential biomarkers were reported in only single studies.^18^ These observations underscore the critical need for larger, multi-cohort studies and robust validation pipelines to enhance the clinical utility and reproducibility of metabolomics findings.

The temporal analysis conducted using our embedding framework further enhances the value of this work. By color-coding publications based on the year of publication and dividing the dataset into discrete time periods, we were able to observe the evolution of research clusters over time (**Figure S3**). Early work in the field was predominantly concentrated in analytical methods (**Figure 2b, Table S4**), while more recent years show a diversification that includes translational research and clinical applications. For example, the cluster related to COVID-19 or SARS-CoV-2 metabolomics^19^ illustrates how the field can rapidly pivot in response to global health challenges, resulting in a densely populated area within the map (**Figure 5b & S5**). Such temporal insights afforded by this technique can be invaluable for policymakers, funding agencies, and research leaders seeking to understand the progression of scientific inquiry and to identify opportunities for interdisciplinary collaboration.

Another noteworthy observation is the impact of publication venues on the dissemination of metabolomics research. Our publication analytics reveal that open-access journals such as Scientific Reports, Metabolites, and PLOS ONE collectively account for close to 50% of all metabolomics publications (**Figure 1c**). The dominance of these platforms underscores the field’s preference for open-access dissemination, which not only facilitates broader reach and collaboration but also accelerates the translation of research discoveries into practical applications.

While our approach has yielded valuable insights, incorporating additional metadata such as full-text analysis, author affiliations, or funding sources could further refine our classifications and provide a more granular view of the research landscape. Looking ahead, the global mapping framework presented here holds significant promise for application in other fast-evolving sub-disciplinary areas. Furthermore, the development of interactive platforms, such as our web application (https://metascape.streamlit.app/), empowers researchers, clinicians, and policymakers interested in the field of metabolomics to explore trends, and identify emerging areas of interest with unprecedented ease.

## METHODS

### PubMed Data Retrieval

We compiled a corpus of PubMed articles by querying for the terms **“metabolomics”** or **“metabonomics”**. Preliminary searches spanned 1993 to early 2024, but as no relevant results appeared before 1998, we restricted our final dataset to the years 1998–early 2024. Data were acquired via the Entrez Programming Utilities (E-Utilities) from the National Center for Biotechnology Information (NCBI), which provide structured XML output. To efficiently manage queries and circumvent the 10,000-record limit of the standard E-Utilities API, we employed the Entrez Direct (E-Direct) package. Two primary E-Direct commands, esearch and efetch, facilitated both the initial search and full-record downloading.

Specifically, we executed:

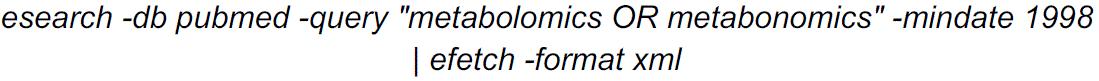

This search looked for the selected terms in various fields of PubMed records, such as: titles, abstracts (summaries), MeSH (Medical Subject Headings) terms, and author keywords. The operation produced an XML file containing over 80,000 entries. We extracted key fields—PubMed ID, article title, abstract text, language, publication year, and author names—using a custom Python script. Records missing abstract text, publication year, or last author name were discarded. For simplicity, we concatenated first and last author names into a single field. The final curated dataset consisted of **80,656** publications, stored in a dataframe for subsequent analyses.

### Publication Classification

To classify publications by research domain, we used journal titles as proxies to infer the primary focus of each article.^6^ We developed a comprehensive keyword mapping spanning 18 predefined categories: oncology, plant biology, nephrology, endocrinology, microbiology, analytical chemistry, pharmacology, neuroscience, food science and nutrition, toxicology, environmental science, animal science, sports science and medicine, epidemiology and public health, developmental biology, aging and gerontology, immunology and vaccine research, and computational biology. For example, “Oncology” included terms such as “cancer,” “oncogene,” and “tumor,” whereas “Microbiology” included such terms as “bacteriology,” “yeast,” and “microbiome.” To standardize classification, journal titles were converted to lowercase, and publications were assigned to a category if the journal title contained any of the category’s keywords. Journals that did not match any category-specific keywords were labeled as “unclassified.” While this approach offered a broad overview of key domains, it resulted in 41,721 publications falling under “unclassified,” reflecting the prevalence of multidisciplinary or nomenclature-agnostic journals in metabolomics. All keyword mappings are provided in **Appendix A**.

### Analysis of Publication Statistics

For each abstract retrieved, the collection of publication metadata such as publication years and journal titles allowed for the assessment of growth trajectories and publication characteristics over time. Abstract length distribution was computed by counting the frequency of both word and character counts. We identified the top publishing journals by counting the number of metabolomics papers published in each journal, highlighting the most prominent venues for disseminating research in this field. Using word cloud visualization with the WordCloud python library (version 1.9.3), we highlighted the most commonly occurring terms across all abstracts. The default stop words in the library were removed in addition to custom stop words ‘used’ and ‘study’’. For the rate of change in publications, to identify overall trends, we computed the year-over-year change rates in publication numbers by calculating the difference in publication counts between consecutive years. To identify periods of consistently low growth and high rates of change in publication trends, we analyzed the annual derivative of publication counts. Using the computed year-over-year change rates, the mean and standard deviation of rates of change for each starting year (Year1) were calculated. The overall mean rate of change across all year pairs was 447.58 publications per year, with standard deviations varying by year. Thresholds were set as follows: (a) High Growth Threshold: Mean rate of change plus one standard deviation for a given Year1, and (b) Low Growth Threshold: Mean rate of change minus one standard deviation for a given Year1. Based on this criterion, years where the mean rate of change exceeded the high growth threshold were classified as periods of consistently high growth. These classifications provided insights into the temporal dynamics of research activity in the field. Early 2024 publications were removed from this analysis.

### Embeddings Generation

Embeddings for publication abstracts were generated using the PubMedBERT model^7^ (microsoft/BiomedNLP-PubMedBERT-base-uncased-abstract-fulltext) *via* the Hugging Face Transformers library (version 4.38.1) and implemented in PyTorch (version 2.1.0+cu121). Each abstract was first preprocessed using the model’s associated tokenizer, which tokenized the text with padding and truncation applied to a maximum input length of 512 tokens. A custom function was implemented to pass these tokenized inputs to the model on a GPU-enabled environment (NVIDIA Tesla T4, CUDA version 12.2 via Google Colab), where the model’s forward pass produced token-level representations. For each abstract, the final 768-dimensional embedding was computed by taking the mean of the last hidden states across all tokens, and the resulting embeddings for all 80,656 publications were stored in a DataFrame as an HDF5 file for downstream analyses.

### Embeddings Dimensionality Reduction

To visualize the higher-dimensional embeddings, we employed two widely used dimensionality reduction techniques: t-Distributed Stochastic Neighbor Embedding (t-SNE) and Uniform Manifold Approximation and Projection (UMAP). Both methods reduced the original 768-dimensional embeddings to two dimensions to facilitate visualization and analysis. The scikit-learn implementation (version 1.2.2) of t-SNE was utilized with the following parameters: n_components = 2 to project embeddings into a two-dimensional space and random_state = 42 to ensure reproducibility. Similarly, UMAP was implemented using the umap-learn library (version 0.5.5) with the same parameters. To evaluate the effectiveness of each dimensionality reduction technique, we used two metrics: k-Nearest Neighbors (k-NN) accuracy and k-NN recall. These metrics provided quantitative insights into how well the reduced embeddings preserved the structure of the original high-dimensional data. The scikit-learn library was used for computing both metrics. k-NN accuracy was determined using the KNeighborsClassifier class. This metric assessed the proportion of correctly classified samples in the reduced space, reflecting how well the two-dimensional embeddings captured the global structure of the data. A k-NN classifier was trained on the two-dimensional embeddings using a train-test split with 1% of the data reserved for testing. The classifier was configured with n_neighbors = 10 (each sample’s label was determined by the majority label of its 10 nearest neighbors), algorithm = brute (to compute distances exhaustively), and n_jobs = –1 (for parallel computations). The accuracy was then evaluated on the test set. k-NN recall measured the local neighborhood preservation between the high-dimensional and reduced-dimensional spaces. For each data point, the k nearest neighbors were computed in both the original high-dimensional space and the reduced 2D space using Euclidean distance. The common neighbors between these two sets were identified by calculating the intersection for each data point. The total number of shared neighbors was then summed across all data points. Finally, the kNN recall was determined by dividing this sum by the total number of possible neighbor comparisons (number of data points multiplied by 10), yielding a measure of neighborhood preservation. This dual-metric evaluation allowed us to quantitatively compare the performance of t-SNE and UMAP, ensuring a rigorous assessment of their ability to preserve the structural characteristics of the high-dimensional embeddings.

### Topic Modelling with a Large Language Model

We performed topic modeling on metabolomics abstracts using a pipeline that combined precomputed PubMedBERT embeddings, dimensionality reduction, and clustering with BERTopic^8^ to identify and represent topics. To streamline the analysis, precomputed embeddings were integrated into the BERTopic pipeline *via* a custom PrecomputedEmbeddings class, which enabled seamless processing without redundant embedding computations. Clustering was conducted with HDBSCAN, an algorithm well-suited for identifying clusters of varying densities and labeling outliers. The hyperparameters were chosen to enhance cluster robustness, with a minimum cluster size of 500 and a minimum sample size of 300. Euclidean distance was employed as the metric for proximity measurement, and the “excess of mass” method was used for optimal cluster selection. Text within each cluster was vectorized using a CountVectorizer, configured to remove common English stop words and capture terms appearing in at least ten abstracts. Both unigrams and bigrams were included to ensure topic specificity. Cluster representations were generated using BERTopic’s class-based Term Frequency–Inverse Document Frequency (c-TF-IDF) approach, which allowed for distinct and interpretable topic descriptions (**Figure 3**). To refine topic labels, we employed OpenAI’s GPT-4o-mini model through BERTopic’s integrated TextGeneration module. Representative documents and c-TF-IDF keywords from each cluster were provided as inputs to generate concise and informative labels. The labeling process was configured to summarize up to 100 representative abstracts per topic while ensuring API efficiency through built-in delays. A prompt consisting of a system prompt, an in-context example, and a main prompt was used (See **Appendix B**) This approach significantly enhanced the interpretability of topics without necessitating manual review of all abstracts. The BERTopic model was trained using our precomputed embeddings and HDBSCAN-generated clusters, with final outputs including topic assignments, probabilities, and refined labels. The pipeline was executed in a high-memory, GPU-enabled computational environment, leveraging Python-based tools including BERTopic (version 0.16.3), HDBSCAN (version 0.8.29), scikit-learn (version 1.2.2), and OpenAI’s API Client (version 1.51.1).

## Data and Code Availability

The code used for the analysis is available at GitHub: https://github.com/obifarin/metamap

The PubMed dataset and associated embeddings can be accessed at Zenodo: https://doi.org/10.5281/zenodo.15020144.20 The first file is a curated XML dataset used in this study, covering the period from 1998 to early 2024. To ensure data quality, entries with errors have been removed. The second file contains embeddings and associated datasets within the same timeframe. It includes PubMed metadata such as PMID, title, abstract, language, journal title, publication year, and authors. Additionally, we provide full 768-dimensional embeddings generated using PubMedBERT, along with uMAP and t-SNE embeddings for dimensionality reduction and visualization.

## Acknowledgements

FMF and OOB acknowledge support by NIH grants 1R01CA218664, 5R61CA281667, R01DK132369, U01DK134191 and the CMaT NSF Research Center (EEC-1648035).

## SUPPLEMENTARY MATERIALS

**Figure S1.**
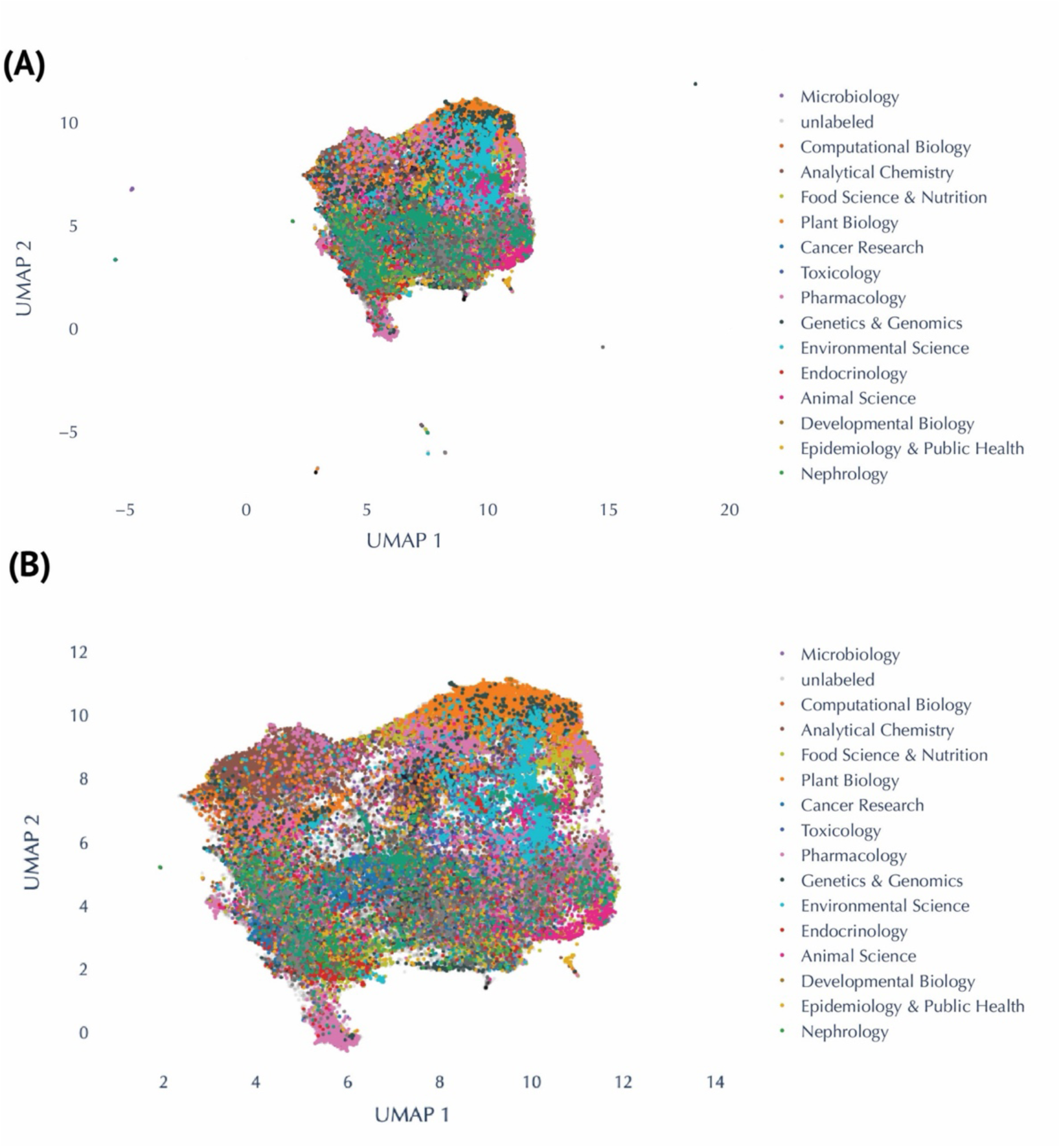
Global visualization of metabolomics research fields using UMAP embeddings. (**A**) Two dimensional UMAP projection of 80,656 publications. (**B**) A magnified view of two dimensional UMAP projection of 80,656 publications.

**Figure S2.**
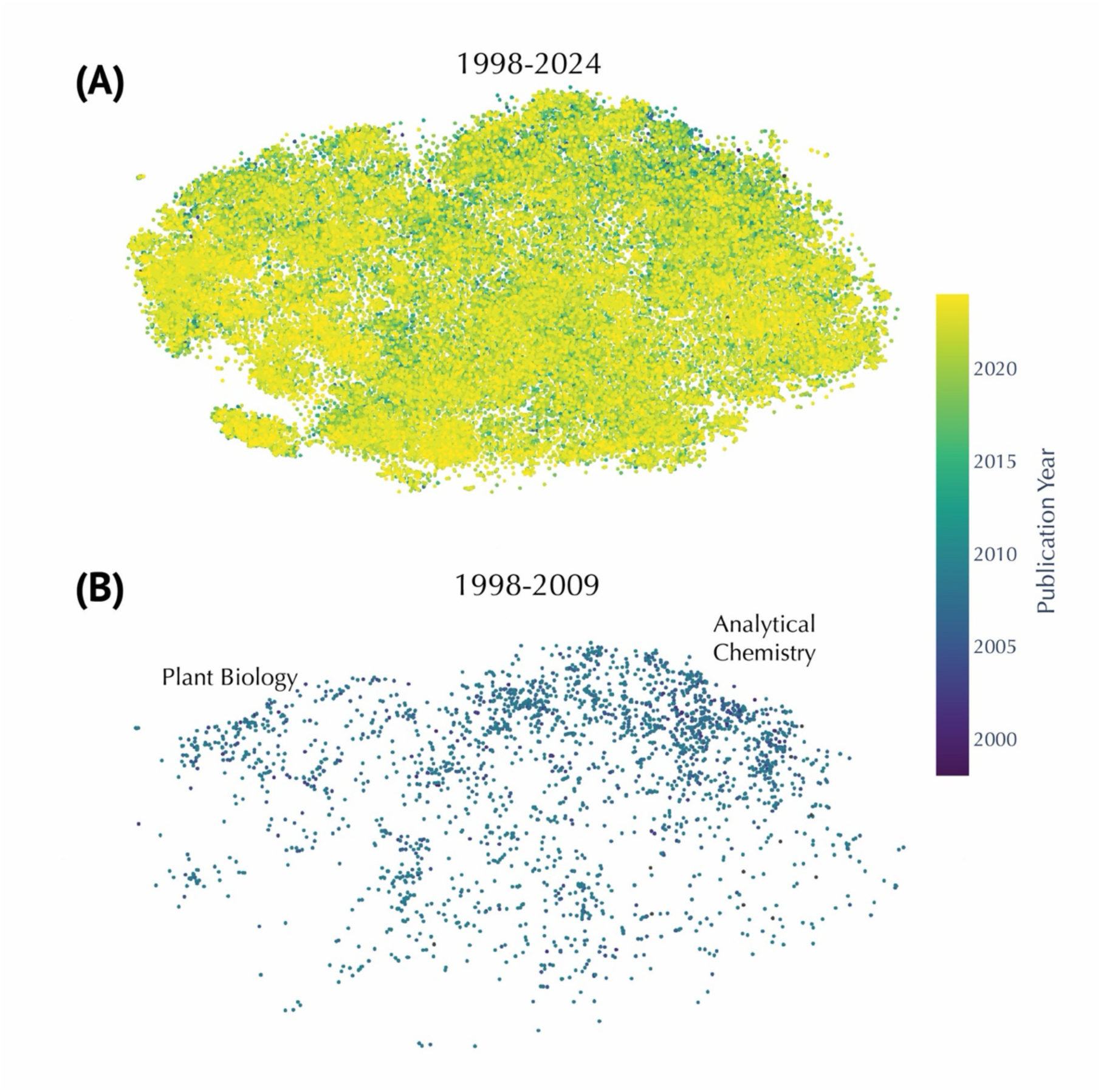
Temporal distribution of metabolomics publications visualized using t-SNE embeddings. (**A**) Showing all publications from 1998 to early 2024. (**B**) Showing all publications between 1998 and 2009. The publications are concentrated in the ‘Analytical Chemistry’ and ‘Plant Biology’ area of the map.

**Figure S3:**
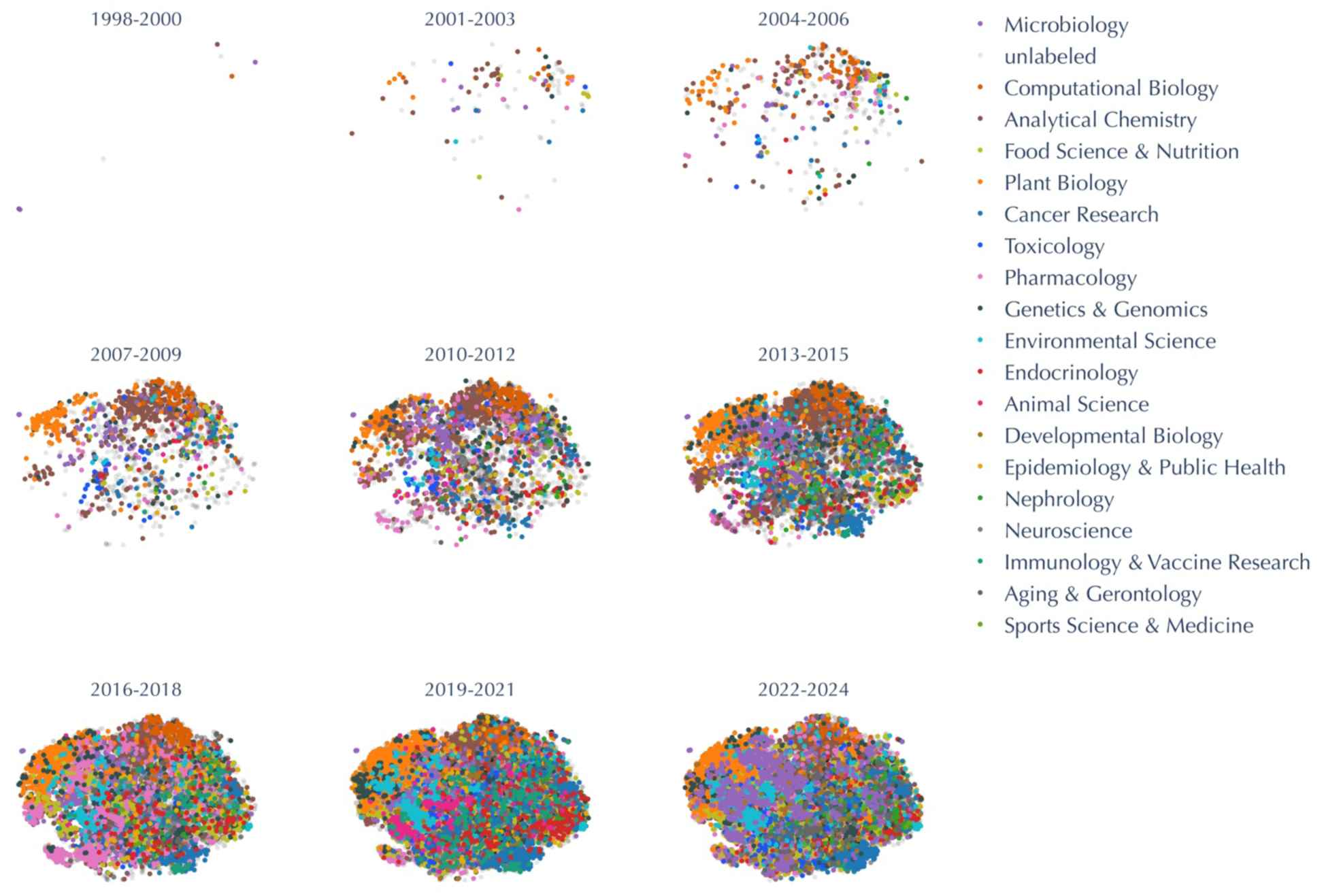
Time-segmented t-SNE visualizations of metabolomics research fields. Clusters represent metabolomics publications divided into eight time periods (1998–2000, 2001–2003, 2004–2006, 2007–2009, 2010–2012, 2013–2015, 2016–2018, 2019–2021, and 2022–early 2024). Each point represents a publication, color-coded by its research domain.

**Figure S4:**
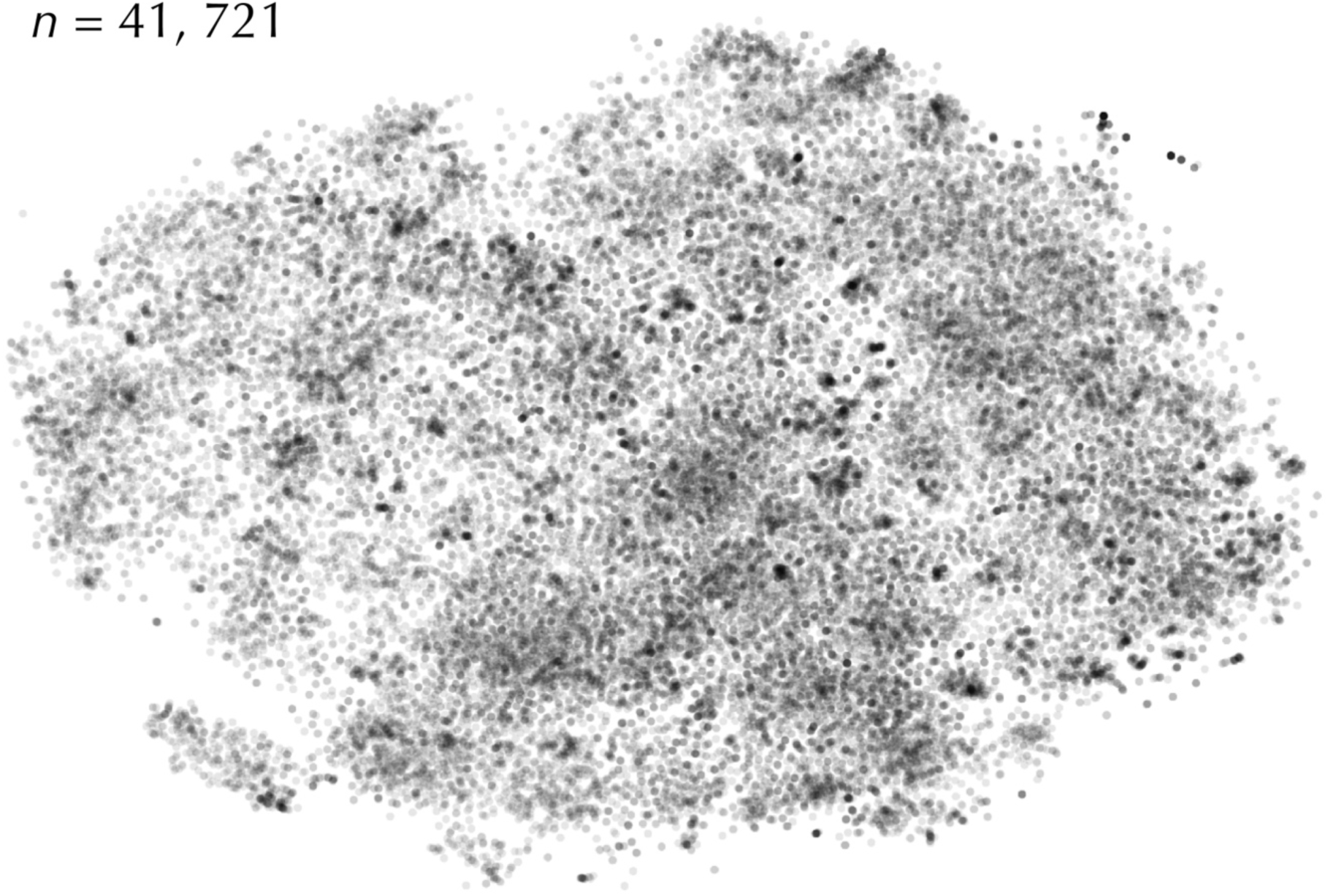
Unlabeled Cluster of Metabolomics Publications. Scatterplot of 41,721 “unlabeled” publications in a t-SNE projection, representing articles not assigned to predefined journal-based categories.

**Figure S5:**
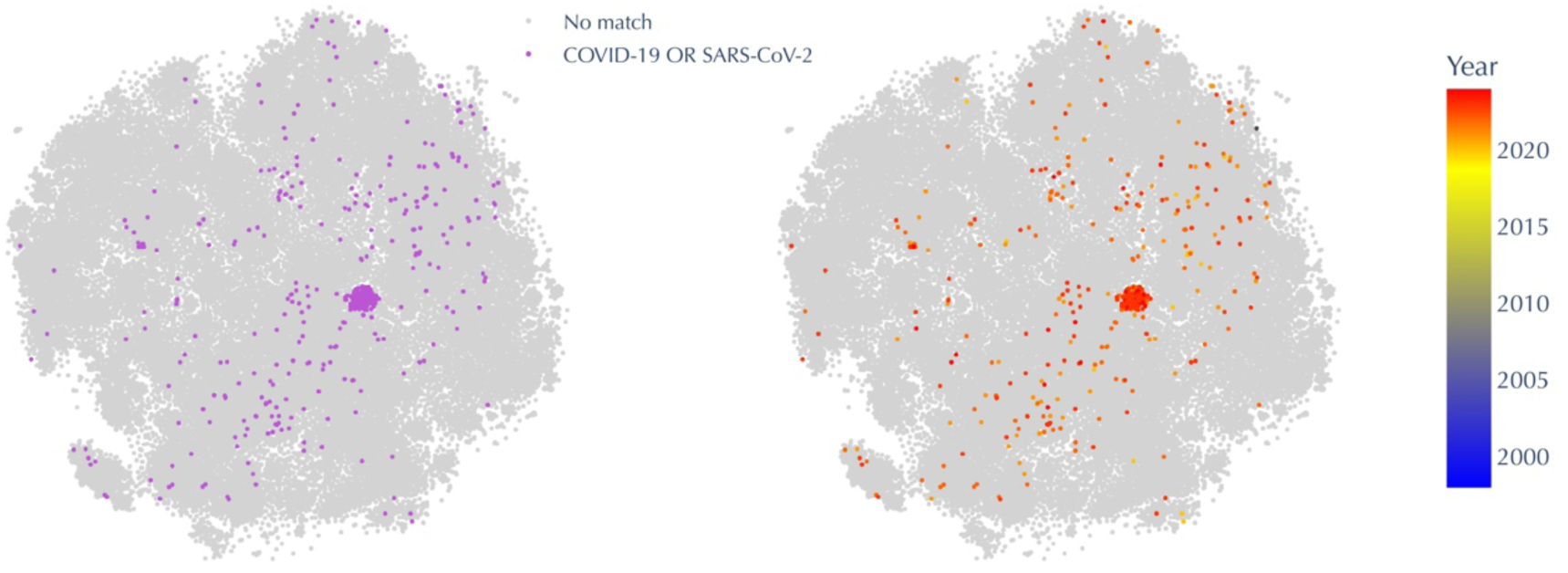
Embeddings illustrating the impact of COVID-19–related research. **Left:** Points in **purple** indicate abstracts mentioning “COVID-19 or SARS-CoV-2,” while **gray** points show no mention. **Right:** The same embedding colored by publication year, transitioning from **blue** (earlier) to **red** (more recent). The visible cluster of purple/red points emphasizes the surge of COVID-19 metabolomics studies in recent years.

**Figure S6:**
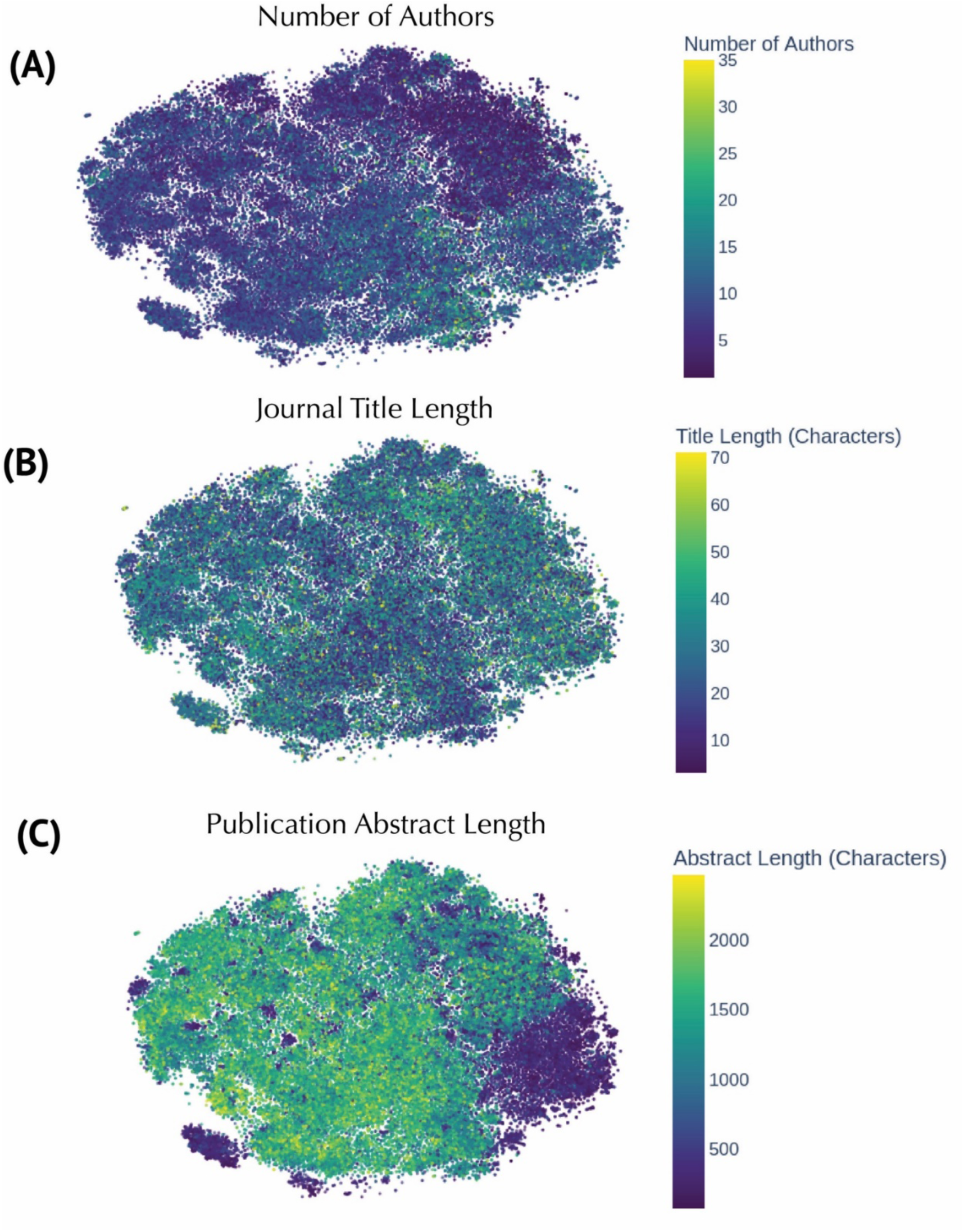
Embeddings colored by publication metadata features. (**A**) t-SNE embeddings of metabolomics publications colored by the number of authors per publication, highlighting collaboration trends across the corpus. (**B**) Embeddings colored by the length of journal titles. (**C**) Embeddings colored by abstract length.

**Table S1:**
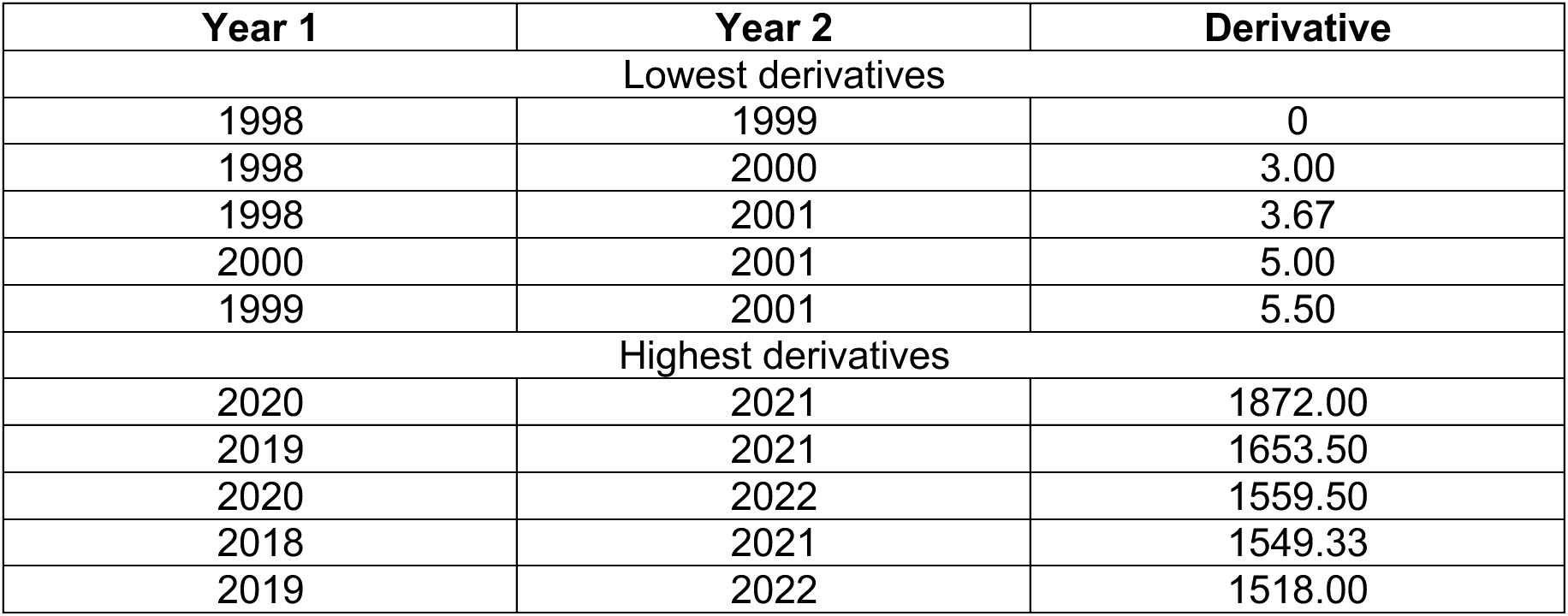
Derivative Analysis of Publication Trends in Metabolomics Research. Yearly derivatives were computed to identify periods of growth acceleration and deceleration. The lowest derivatives occurred during the nascent stages of the field (1998–2001), while the highest derivatives were observed during recent years (2019–2022).

| Year 1 | Year 2 | Derivative |
| --- | --- | --- |
| Lowest derivatives |  |  |
| 1998 | 1999 | 0 |
| 1998 | 2000 | 3.00 |
| 1998 | 2001 | 3.67 |
| 2000 | 2001 | 5.00 |
| 1999 | 2001 | 5.50 |
| Highest derivatives |  |  |
| 2020 | 2021 | 1872.00 |
| 2019 | 2021 | 1653.50 |
| 2020 | 2022 | 1559.50 |
| 2018 | 2021 | 1549.33 |
| 2019 | 2022 | 1518.00 |

**Table S2:** Years with Consistently High and Low Rates of Publication Change in Metabolomics Research. Mean and standard deviation (Std) rates of change were calculated for each starting year (Year1). High-growth years (e.g., 2011–2021) were characterized by substantial adoption of metabolomics techniques, while low-growth years (e.g., 1998–2002) reflect the early development of the field. Thresholds for high and low growth were defined as the mean rate of change ± one standard deviation.

| Year1 | Mean | Std |
| --- | --- | --- |
| Years with consistently high rate of change |  |  |
| 2011 | 627.57 | 179.10 |
| 2013 | 766.47 | 176.62 |
| 2014 | 796.21 | 214.89 |
| 2015 | 924.82 | 186.25 |
| 2016 | 942.88 | 264.33 |
| 2017 | 1151.25 | 217.46 |
| 2018 | 1412.38 | 98.406 |
| 2019 | 1477.12 | 147.45 |
| 2020 | 1563.06 | 307.18 |
| 2021 | 1098.75 | 209.66 |
| Year1 | Mean | Std |
| Years with consistently low rate of change |  |  |
| 1998 | 159.94 | 155.85 |
| 1999 | 176.80 | 161.29 |
| 2000 | 195.68 | 166.85 |
| 2001 | 218.96 | 170.63 |
| 2002 | 242.70 | 175.83 |

**Table S3:** Publication counts by research fields in metabolomics. Breakdown of all publications across 18 predefined research domains, based on journal titles. The “unlabeled” category accounts for publications without domain classification.

| Category | Count |
| --- | --- |
| unlabeled | 41721 |
| Analytical Chemistry | 8981 |
| Plant Biology | 4648 |
| Pharmacology | 4229 |
| Food Science & Nutrition | 4053 |
| Microbiology | 3605 |
| Cancer Research | 2657 |
| Environmental Science | 1945 |
| Genetics & Genomics | 1312 |
| Toxicology | 1175 |
| Neuroscience | 980 |
| Endocrinology | 953 |
| Computational Biology | 943 |
| Immunology & Vaccine Research | 924 |
| Animal Science | 876 |
| Epidemiology & Public Health | 547 |
| Aging & Gerontology | 448 |
| Developmental Biology | 389 |
| Nephrology | 277 |
| Sports Science & Medicine | 76 |

**Table S4:** Publication counts by research fields in metabolomics from 1998 to 2009.

| **Category** | **Count** |
| --- | --- |
| unlabeled | 1116 |
| Analytical Chemistry | 577 |
| Plant Biology | 205 |
| Pharmacology | 155 |
| Microbiology | 109 |
| Computational Biology | 106 |
| Genetics & Genomics | 82 |
| Food Science & Nutrition | 72 |
| Toxicology | 61 |
| Cancer Research | 45 |
| Endocrinology | 21 |
| Environmental Science | 20 |
| Neuroscience | 18 |
| Nephrology | 9 |
| Animal Science | 8 |
| Epidemiology & Public Health | 7 |
| Developmental Biology | 6 |
| Immunology & Vaccine Research | 6 |
| Aging & Gerontology | 2 |

**Table S5.** Publication counts by topics in metabolomics. Breakdown of all publications across the topics generated by GPT4o mini and the BERTopic pipeline. The “outlier” category accounts for publications without topic assignment.

| Custom Name | Count |
| --- | --- |
| Outlier | 14980 |
| Plant Stress Response Mechanisms | 19258 |
| Metabolic Profiles and Dysregulation | 9730 |
| Cancer Metabolism and Therapy Resistance | 7254 |
| Metabolomics Data Analysis and Integration | 6499 |
| Gut Microbiota and Metabolomic Interactions | 4313 |
| Metabolomics in Neurodegenerative Disorders | 2723 |
| Environmental Toxicology and Metabolism | 2389 |
| Metabolomics in Animal Nutrition | 1971 |
| Microbiota-Gut-Brain Axis Interactions | 1661 |
| Kidney Disease Metabolomics and Biomarkers | 1479 |
| Metabolomics in Rheumatoid Arthritis | 1318 |
| Maternal Metabolomic Changes in Pregnancy | 1238 |
| Lung Disease and Metabolic Dynamics | 1186 |
| NMR Spectroscopy Innovations in Metabolomics | 779 |
| COVID-19 Metabolomic and Immune Responses | 776 |
| Lipid Profiling Techniques | 687 |
| Host-Parasite Metabolic Interactions | 656 |
| Male Fertility and Reproductive Metabolomics | 652 |
| Ocular Metabolomics and Disease Mechanisms | 604 |
| Salivary Metabolomics in Oral Health | 503 |

## Appendix A: Code Showing Label Mapping for Label Inference using Journal Titles

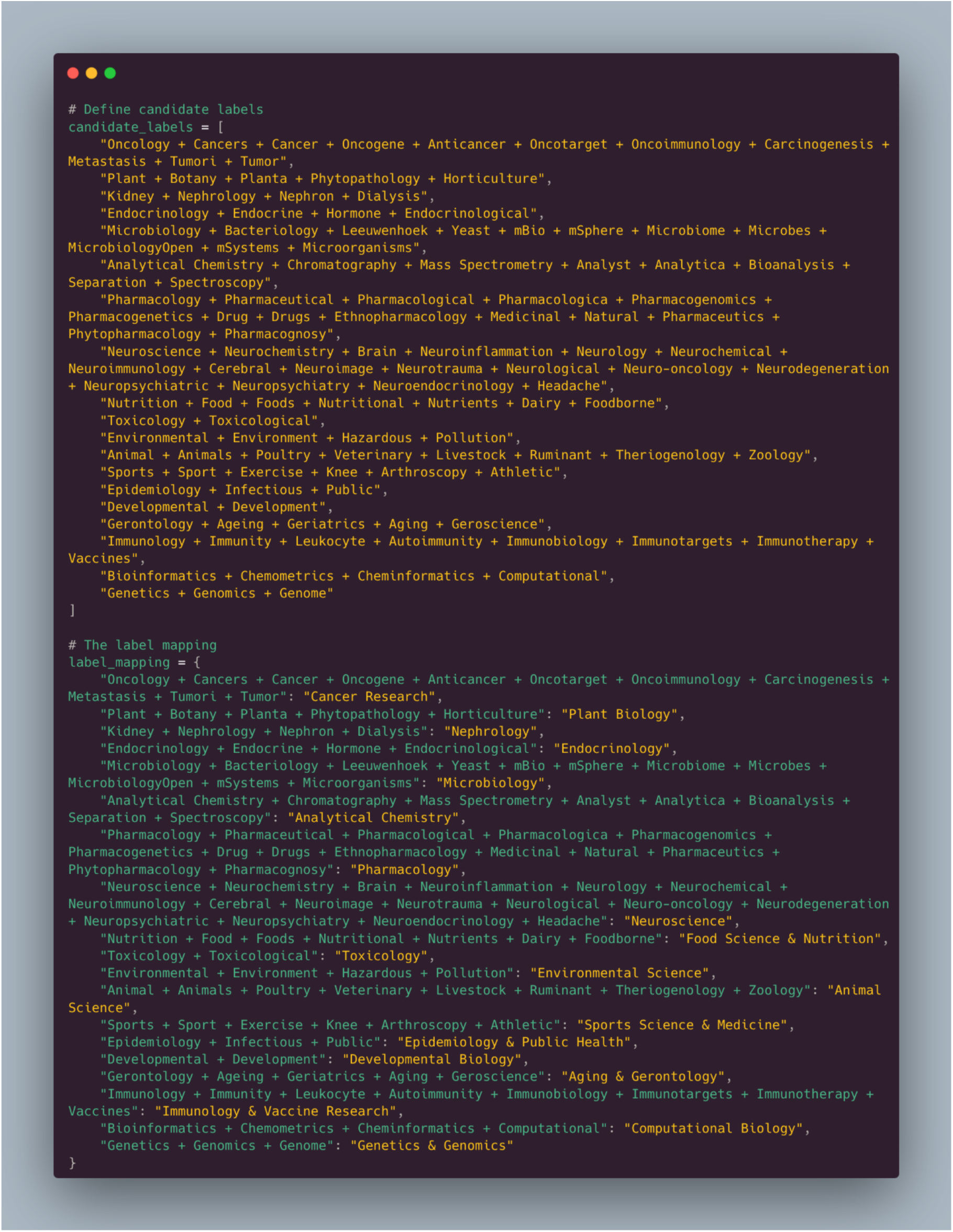

## Appendix B: Prompt for Topic Modelling

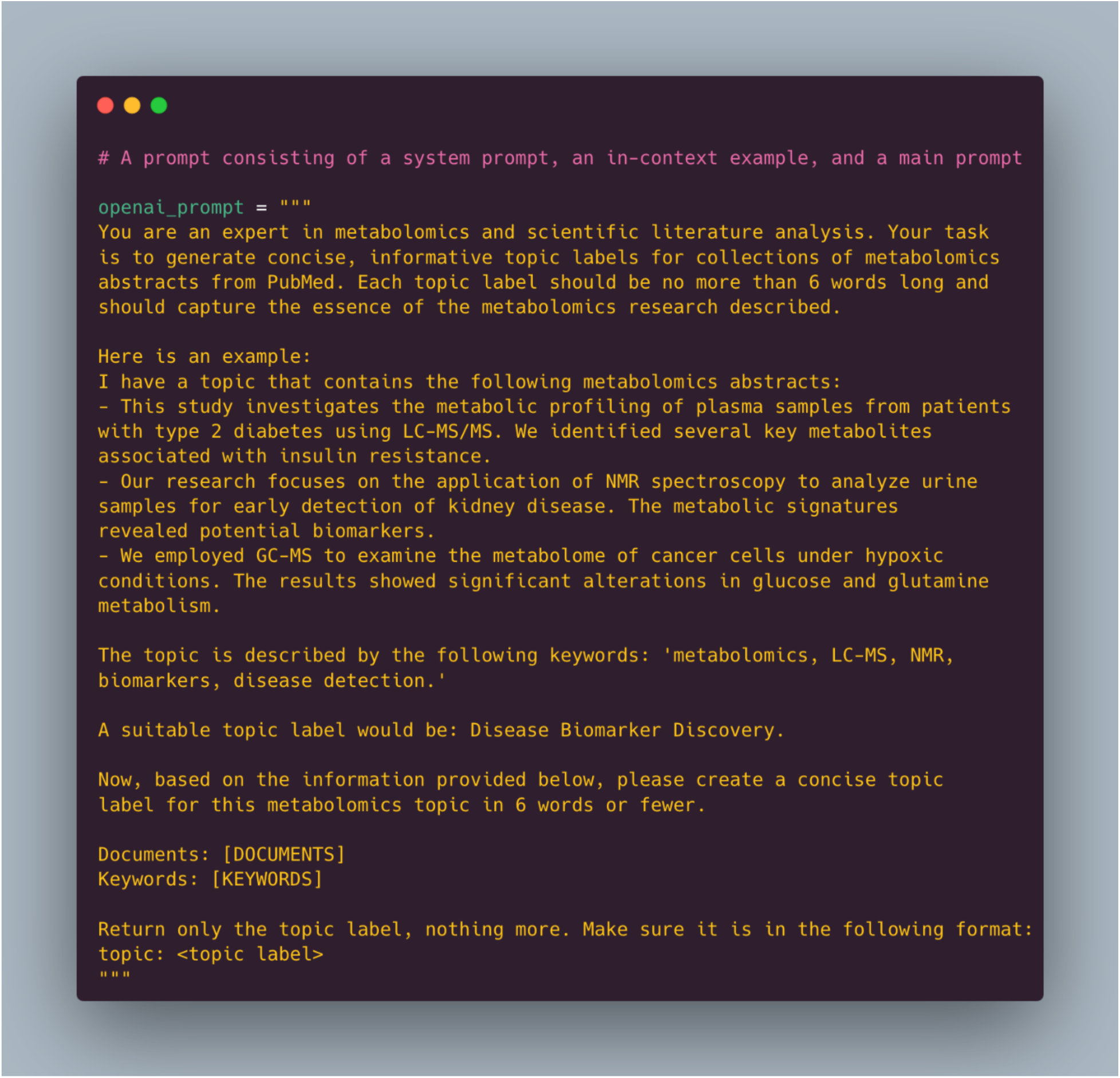

